# Prediction of Polygenic Risk Score by Machine Learning and Deep Learning Methods in Genome-wide Association Studies

**DOI:** 10.1101/2022.12.30.522280

**Authors:** R. Onur Öztornaci, Erdal Coşgun, Cemil Çolak, Bahar Taşdelen

## Abstract

Polygenic risk score (PRS) is a method that using multiple SNPs simultaneously and can be calculated as a typical disease risk score. It is useful method for precision and personalised medicine. Calculating PRS with the classical method, it is frequently used to use two different data sets which are training and testing sets. It is a disadvantage for the classical method. By using a single data set, machine learning (ML) and deep learning (DL) methods both avoid the problem of overfitting and can be used as a good alternative method. Genome-wide Association Studies (GWAS) data were generated with the PLINK Program by replicating a hundred times at different allele frequencies and different sample size. We applied two different ML algorithms which are Support Vector Machine (SVM) and Random Forest (RF) as well as DL approach. ML methods can obtain more consistent results in terms of case-control separation compared to PRS calculated with the classical method (PRS). The use of ML and DL methods as an alternative to classical methods to calculate PRS has been suggested.

## INTRODUCTION

Genome-wide association studies (GWAS) are methods for applying to find out and investigating the genes and genomic regions may have a reason of specific disease. By using whole chromosomes and thus hundreds of thousands of gene simultaneously, gene-gene and gene-environmental relationships are investigated. Therefore, GWAS are not only conducting on specifically selected genes but also aiming determine differences between case and control groups so that using big data sets. Changes in single nucleotides in the genome sequence are called single nucleotide polymorphism (SNP), which explains to a great extent why some individuals are healthier while others are prone to disease, why the same disease progresses differently among different individuals, and why some individuals respond positively to treatment while others do not. GWAS are conducted with SNP data. Nowadays, SNPs that cause many diseases have been identified. Therefore, studies are need far beyond the GWAS. Those studies such as polygenic risk score (PRS), it may help for clinicians and, it would be useful to precision medicine and personalized medicine before disease would not be development [1,2]. PRS is a risk criterion that uses so many SNPs at the same time for calculating a risk score of any genetic disease. A low PRS means that the risk of genetic disease susceptibility is low, a high PRS, on the contrary, means a high risk of predisposition to genetic disease. Aims of this study, to find out the model that most accurate to consider to separation of case-control and predicting genetic risk, besides, validation of classical PRS calculating by using machine learning (ML) and deep learning (DL).

## MATERIALS AND METHODS

In this study content for calculating PRS, as a purpose of finding out the best model of PRS, raw GWAS data set (which is bad, bim, fam files) that has been simulated from 1000 genome project real datasets, it is consisting of 251 case and 232 controls as well as 489805 SNPs that associated with the obesity is used. To determine cases and controls, above the 30 BMI was used as a criterion [3]. Our simulation scenarios, we created different odds ratios for each case and control groups and different sample sizes. If compared with all SNPs, number of the SNPs associated with disease have %1 rate are created for cases groups in all datasets. For the purposes of separating cases and controls group, the SNPs associated with disease have been created by less than 0.05 p-value is obtained from logistic regression analysis.

### Poligenik Risk Score

Polygenic risk score (PRS) is a method useful for calculating a complex genetic diseases risk as well as by nature of PRS, whole variants on genomes are using for calculation and it could be including more information using whole variants on genome than individual mutations. PRS is calculated with the sum of all variants on genomes.

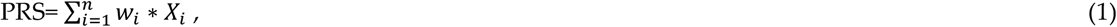

It is calculated by (1) formulation. Where, *w_i_* i-th is weights of SNP, while *X_i_* express the alleles affected by the weights^3^.The odds ratio is used for each weight vector.

### Machine Learning Methods

Machine learning (ML) is a name of the system of algorithms which are catching patterns form the data that used and improving prediction rate by themself. The other word, the ML algorithms are having self-improvement for increasing performance of the prediction by patterns on used datasets [4]. Machine learning are increasingly becoming most popular methods nowadays because compared with classical statistical methods they have no assumption like normality, sample size ect. Besides, having so many tunning parameters when solving nonlinear problems an advantages of ML methods [5,6]. By the nature of genetic epidemiological datasets are big data; therefore, the mathematical methods which are used ought to be able to support this situation as well. Generally, due to ML methods are having good prediction power, it has been become more applying methods when analysing big data^7^.

### Support Vector Machines

Support vector machines (SVM) invented by Vladimir Vapnik in 1992. Lagrange multipliers are using for solving classification problems thus it can be finding the lines among the classes that separated with minimum error rate. Though the training time is higher than the other algorithms if compared, SVM is resistant to over-fitting problem [8,9].

### Random Forest

Random Forest (RF) is an algorithm consisting of combining a lots of decision trees. It can be used for both categorical and continuous data as well as classification and regression models. RF also can be used for missing data imputation, feature selection, finding contribution of variables on model which is call variable importance [10–13].

### Deep Learning Methods

Deep learning (DL) is a branch of the machine learning (ML) however, the differences between ML and DL is processing data such as feature selection, data labelling ect. before the application of the model. The data should have processed before applying ML while DL does not need this [14]. The major advancements have been in image and speech recognition as well as natural language processing and language translation. The successes of deep learning originate from how it learns hierarchical representations of data by increasing the level of abstraction [15,16]. Deep learning architectures are artificial neural networks of multiple non-linear layers in which each successive layer operates on the representation from the preceding layer. Each layer consists of one or more artificial neurons, each of which is connected to other neurons in the preceding layer. The artificial neuron receives separately weighted inputs and sums them to produce an activation using an activation function such as sigmoid and rectified linear unit (ReLU). In the training step, the most suitable hierarchical representations can be learned from data by optimizing the weight parameters in each layer. Once the forward pass sequentially propagates the output function signals forward through the network, in the final output layer, the loss function calculates the error. To minimize the error, the backward pass back-propagates error signals and updates weight parameters using optimization algorithms based on stochastic gradient descent (SGD) [17].

### Alternative Methods for Polygenic Risk Score Prediction

Firstly, we would like to identify weight vectors for developing a new method when predicting PRS. Weight vectors as determined by the *SVMW_i_*. and *DLW_i_* for the SVM and DL methods. In these methods, weight matrix for determining of the classes are used in vector of weights when predicting PRS. *RFI_i_*, variable importance *I_i_* measurements for RF method, was used as weight vector. For all methods, each SNPs multiplying with weight vector and thus, individually risk scores were obtained. The formulas are as follows.

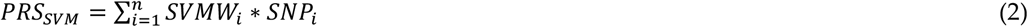

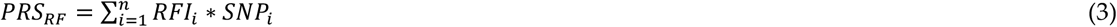

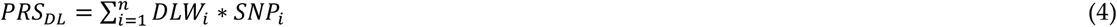

### Simulations Steps

It is very hard and expensive to access to GWAS data for a specific disease, therefore; we simulated data with PLINK software, the raw files (bed, bim, fam) were used for developing a new method as well as comparing to power of approaches. Samples size of the simulations for case-controls: 500-500, 1000-1000, 2000-2000. Number of the SNPs: 5000, 10000, 50000, 100000 are selected. Software and operating systems are PLINK, R, Python, Linux, Microsoft Azure (Figure 1) [18–22].

**Figure 1.**
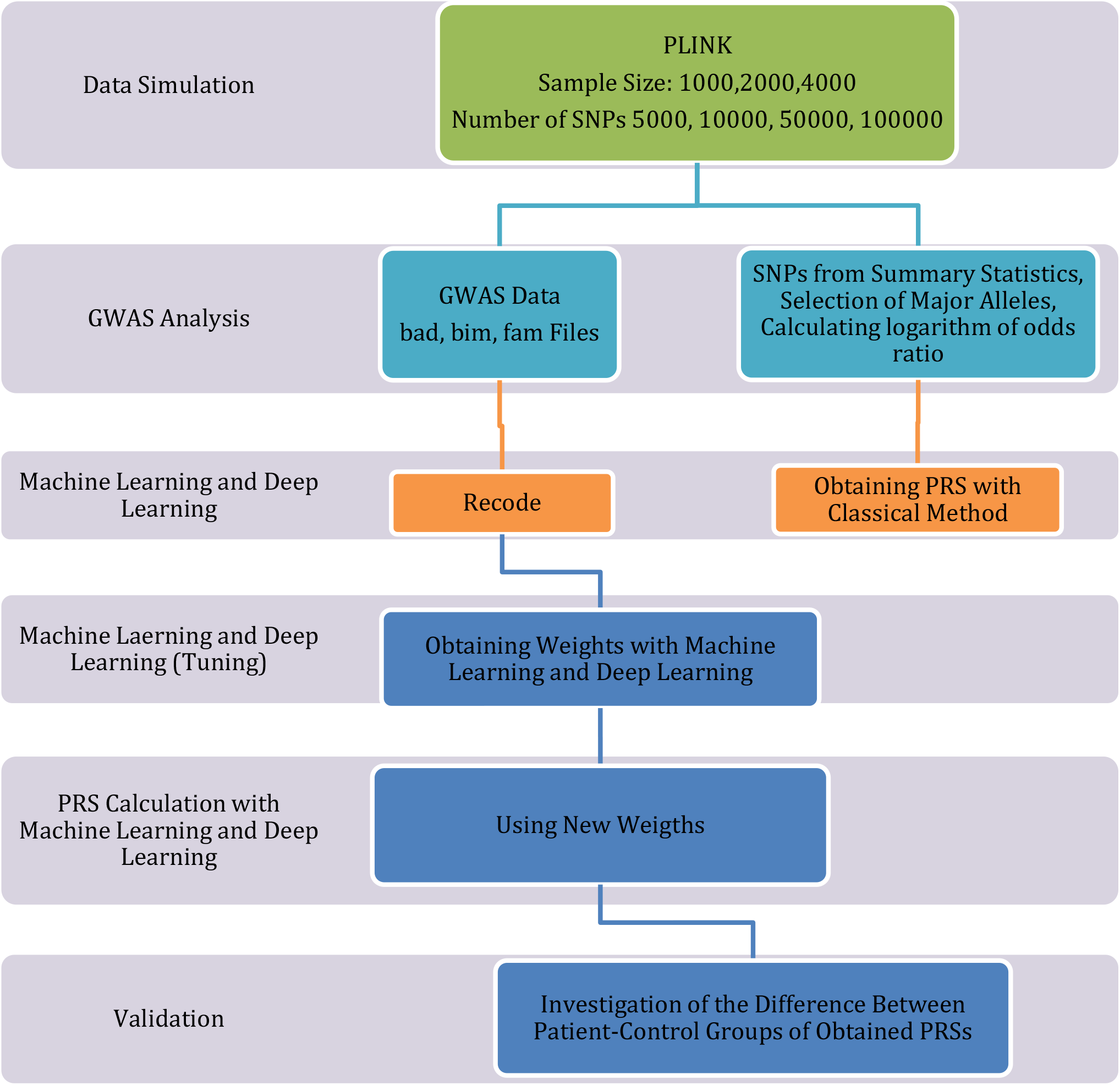
Pipeline for Obtaining a Polygenic Risk Score

## RESULTS

In simulation study, we compared polygenic risk scores calculated by classical method, SVM, RF and DL methods for different SNP numbers and sample sizes. Statistically significant difference was observed between the case-control groups in terms of PRS calculated by the classical method in all SNP numbers and all sample sizes (p<0.001). While the lowest results in both control and patient groups were obtained with the classical method PRS in all sample sizes, it was observed that the highest results were obtained with DL. It was observed that a more homogeneous variation range was obtained in the RF method, the similar interpretations can be said for DVM. It can be said that all newly developed methods can be used as an alternative to the classical method in low SNP numbers. While there was a statistically significant difference in genetic risk score calculated with the classical PRS calculation method DVM and RF for 10,000 SNP between 500 cases with the smallest sample size and 500 control groups (p<0.001), no statistically significant difference was found when the DL method was used (p =0.361). When number of the SNPs is 50000, classical method, SVM and RF methods have statistically differences between cases and control groups regarding PRS (p<0.001), however, DL method does not any differences (p=0.642). When number of the SNPs is 100.000; classical method, SVM and RF can separate to cases and controls in terms of PRS (p<0.001) while DL method is not (p=0.803). Therefore, changing of the number of the SNPs is just important for DL method. When working with small number of SNPs, DL method is best model for separating cases and controls but when number of the SNPs is increased RF, SVM and classical method better than DL can be said. RF and SVM can be used instead of the classical calculation method in all SNP numbers, and it can be said that these methods have narrower confidence intervals and make the case-control separation more successfully as the number of SNPs increases (Table 1).

**Table 1.**
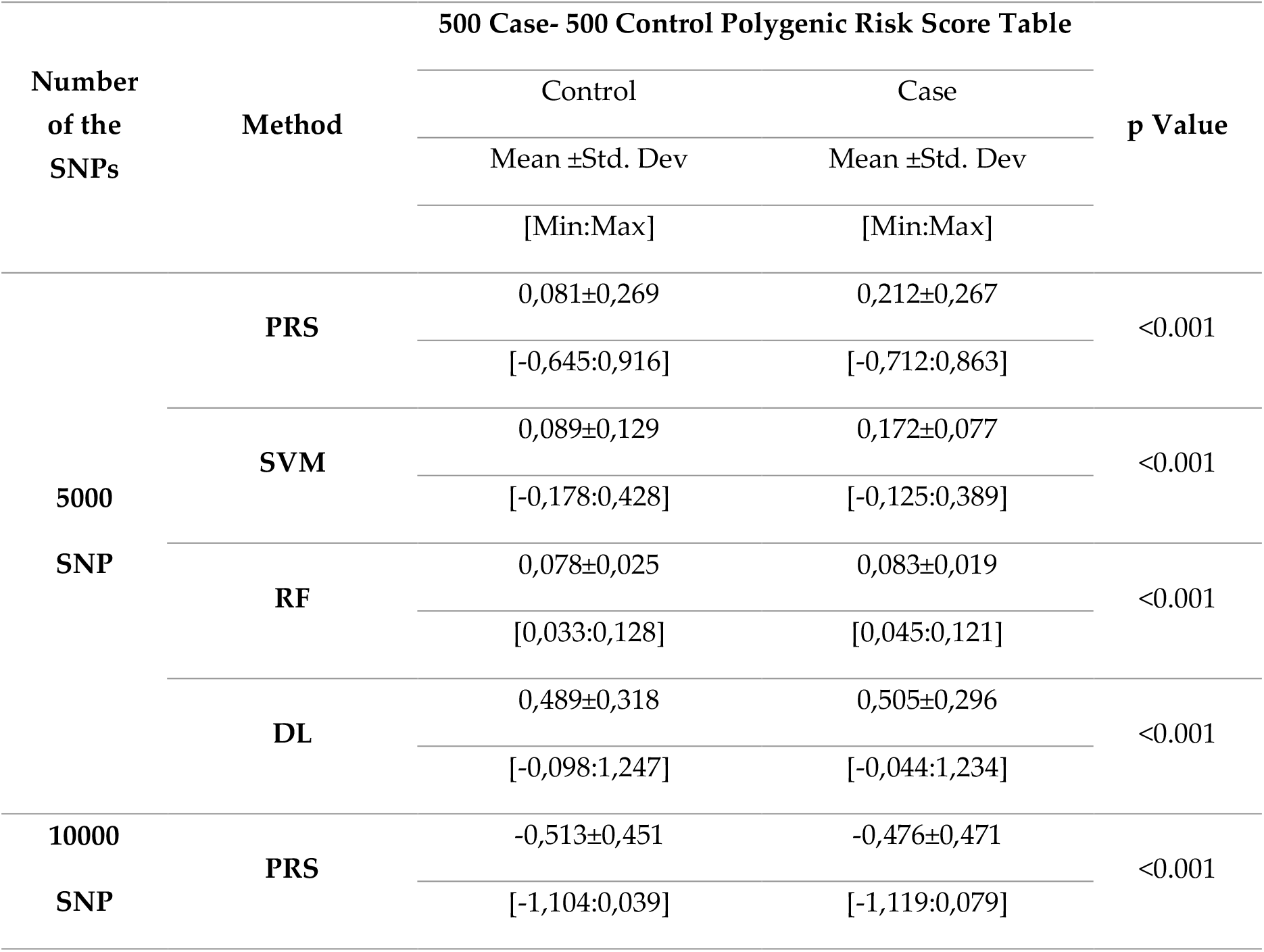
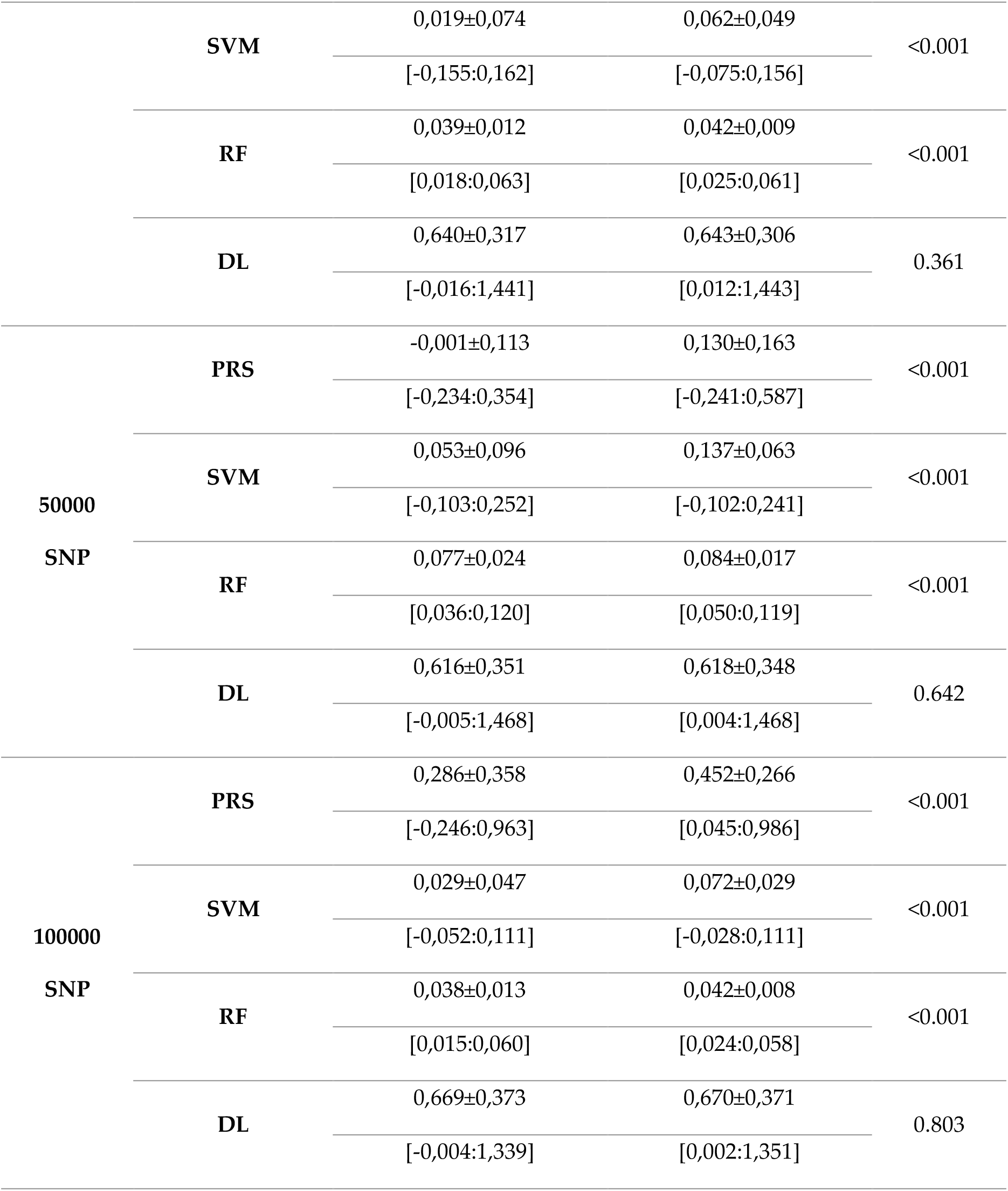
Comparison of Classical Polygenic Risk Score (PRS) and Risk Scores Weighted According to Other Models in Case and Control Groups (N:500-500)

The best and powerful model is SVM for separating case and controls when number of sample size 2000. For SNPs 5000, all methods can separate each other cases and controls so that all methods are useful for predicting PRS; however, for 10000 SNPs, classical method, RF and SVM can be used for predicting to PRS (p<0.001) while DL cannot useful because, it cannot be separated cases and controls (p=0.467). For 50,000 SNPs, there was a statistically significant difference between the case-control groups in terms of genetic risk score calculated with the classical calculation method, DVM and RF (p<0.001), while there was no statistically significant difference when the DL method was used (p=0.652). Similar situation for 100,000 SNPs, while there was a statistically significant difference between the case-control groups in terms of genetic risk score calculated with the classical calculation method, SVM and RF (p<0.001), no statistically significant difference was found when the DL method was used (p=0.825) (Table 2).

**Tablo 2.**
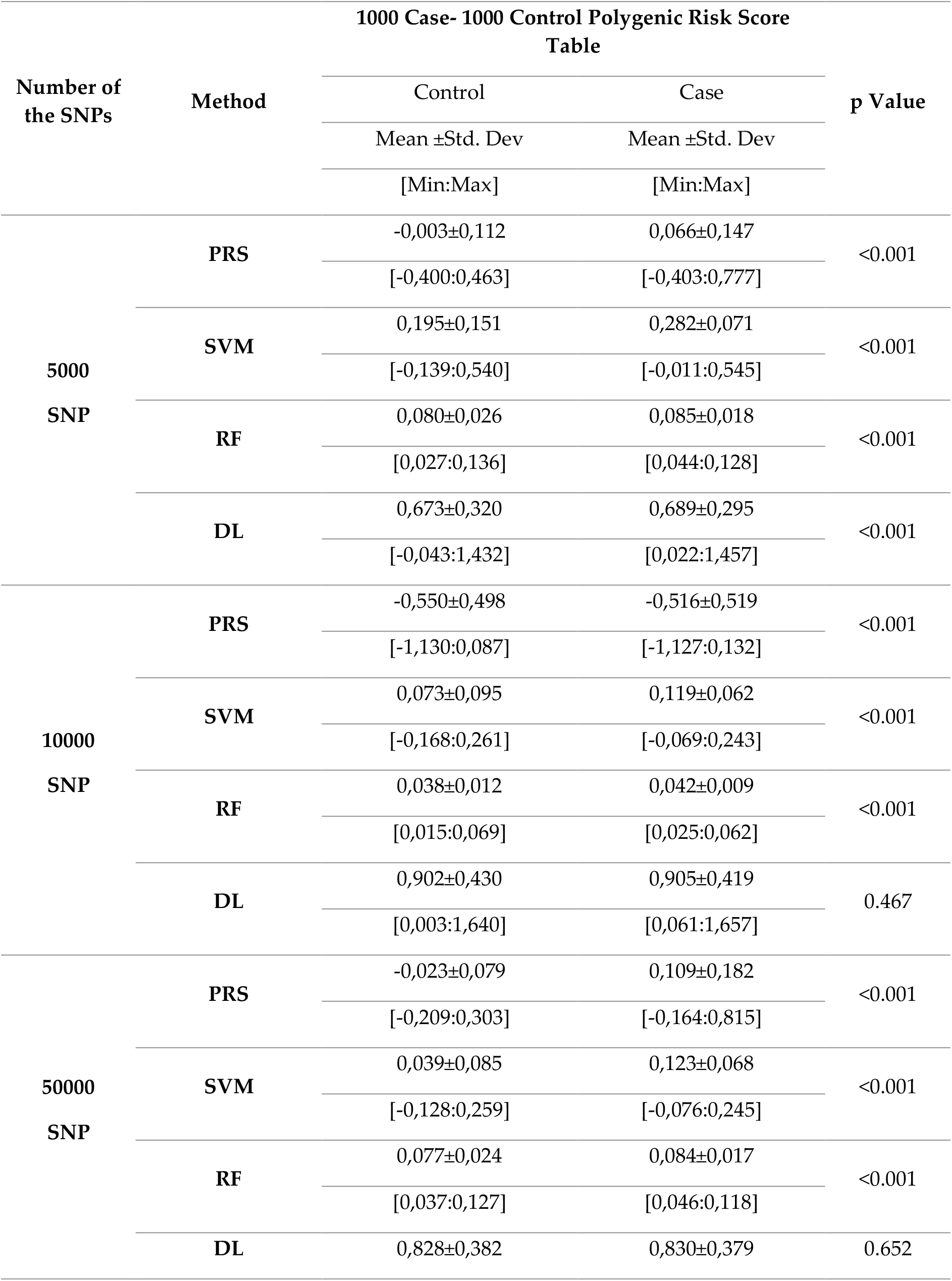
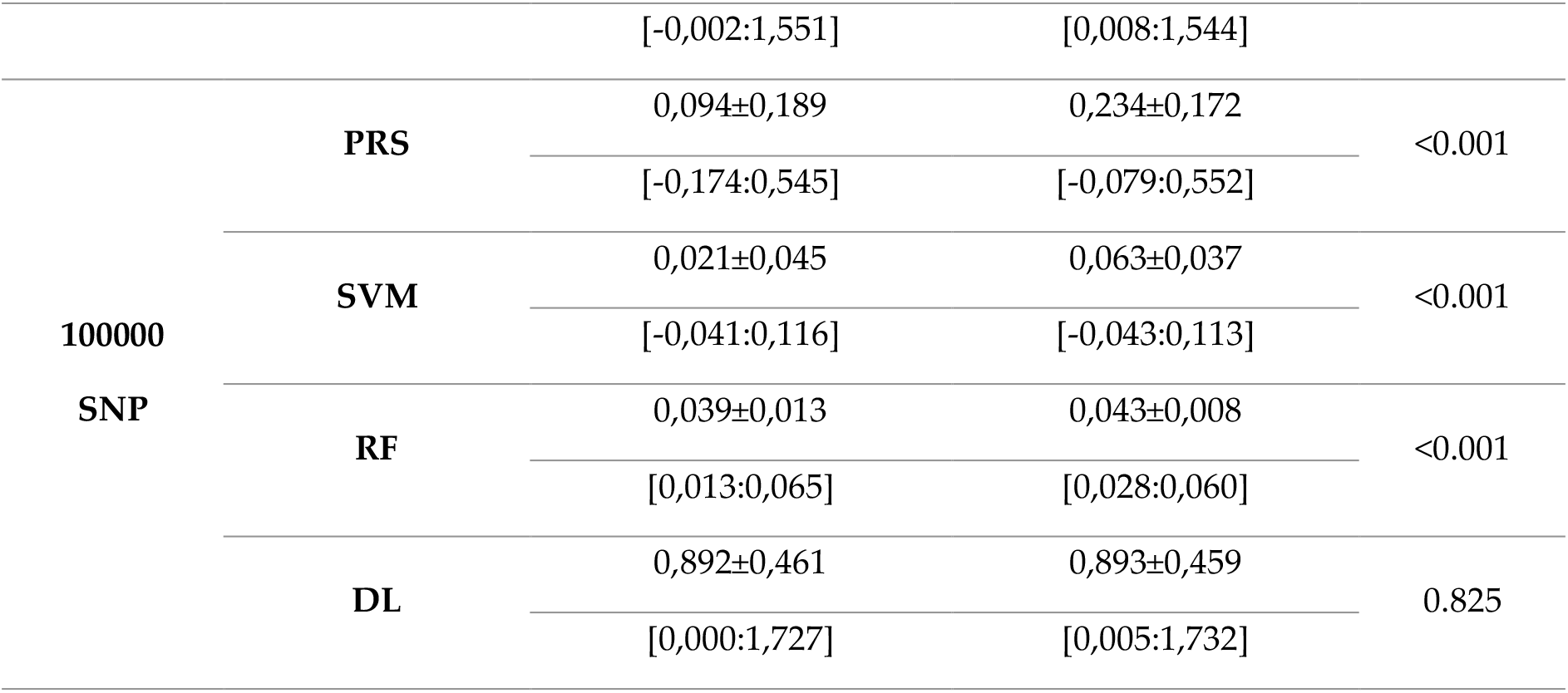
Comparison of Classical Polygenic Risk Score (PRS) and Risk Scores Weighted According to Other Models in Case and Control Groups (N:1000-1000)

For the highest sample size of 2000, the DL method obtained the highest PRS for 5000 TNPs for both case and control groups, followed by the SVM method in the range of [0.20:0.40]. The lowest results PRS were obtained by the classical method. While there was a statistically significant difference between the case-control groups in terms of the PRS calculated with the classical method, DVM and RF (p<0.001), no statistically significant difference was found when the DL method was used (p=0.843). When the number of SNPs is 50000, it is seen that the PRS obtained with classical method give lower results in both control and case groups compared to other methods. RF and SVM methods have obtained results close to each other. While there was a statistically significant difference between the Patient-Control groups in terms of genetic risk score calculated with the classical PRS calculation method DVM and RF (p<0.001), no statistically significant difference was found when the DL method was used (p=0.760). The same situation is seen that when 100000 SNPs was used (Table 3).

**Tablo 3.**
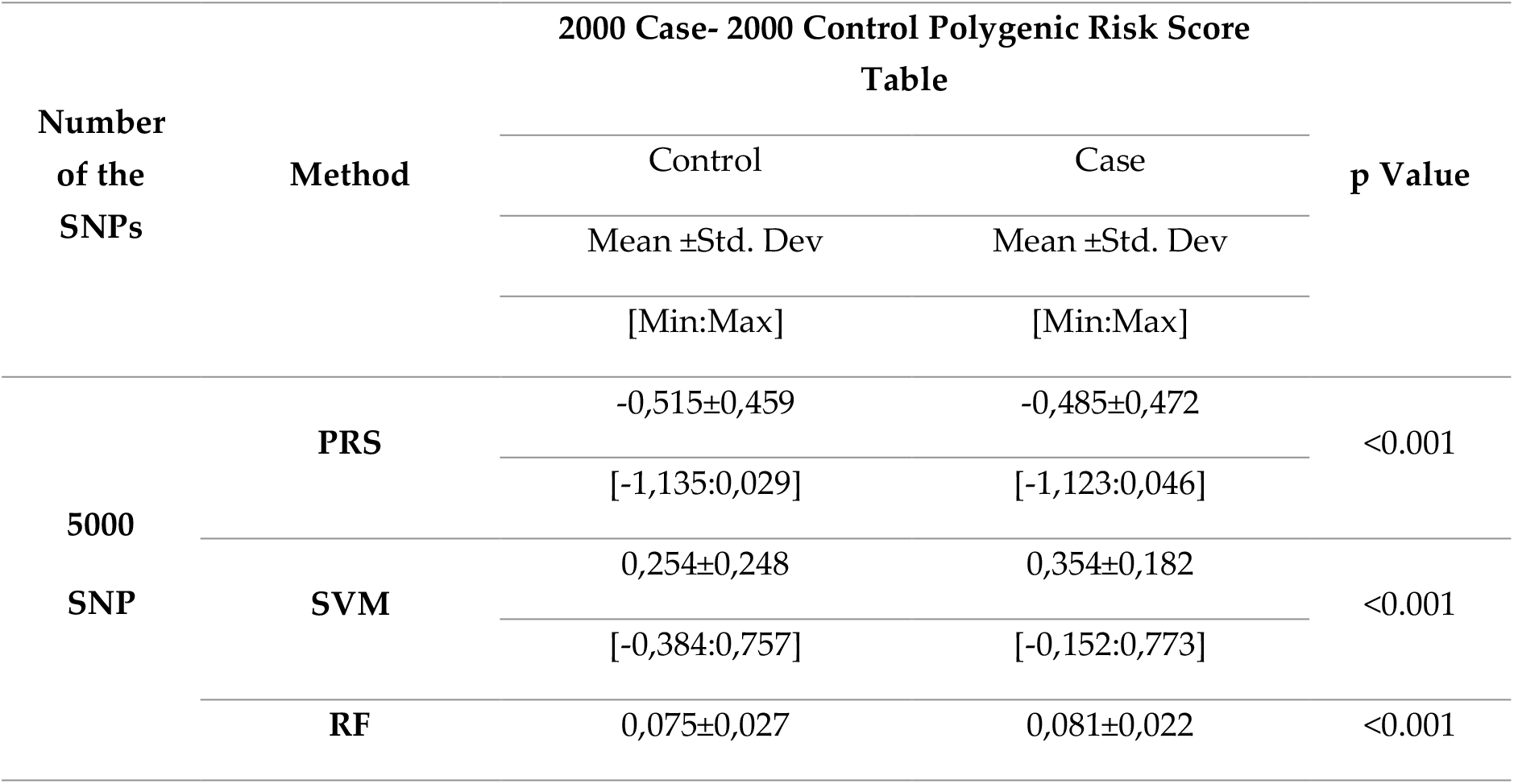
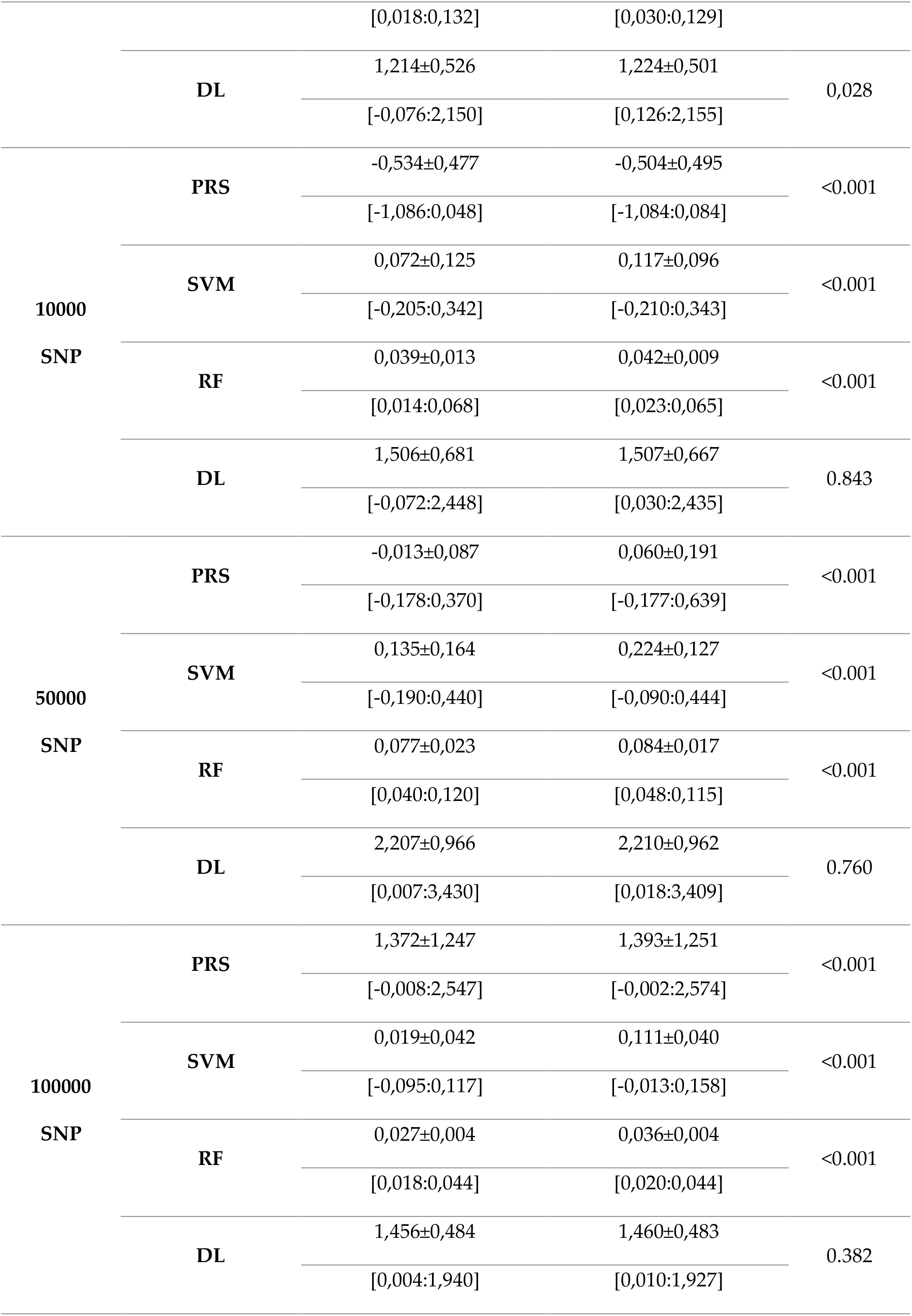
Comparison of Classical Polygenic Risk Score (PRS) and Risk Scores Weighted According to Other Models in Case and Control Groups (N:2000-2000)

While there was a statistically significant difference between the case-control groups in terms of the classical calculation method and the PRS calculated by SVM (p<0.001), no statistically significant difference was found when the DL method (p=0.989) and RF method (p=0.290) were used. If separate evaluations are made for the case and control groups; In the control group, the mean value of −0.214 for the classical PRS calculation method is within the range of [−0.246:0.963] calculated by simulations. Similarly, the mean value of 0.178 for the DL method is within the range of [−0.004:1,339]. For the SVM method, the simulation results are very close to the average values obtained from the real data. However, there is no similarity for the RF method. When investigating case groups; for classical method, DVM and DL method, they include the mean value obtained from real data and [min: max] values obtained because of simulation (Table4).

**Tablo 4.** Comparison of Classical Polygenic Risk Score (PRS) and Risk Scores Weighted by Other Models Using Real Data with Patient Group and Healthy Control Group

| 251 Case- 232 Control Polygenic Risk Score Table |  |  |  |
| --- | --- | --- | --- |
| Method | 489805 SNP |  | p Value |
|  | Control | Case |  |
| | Mean ±Std. Dev | Mean ±Std. Dev | |
|  | [Min:Max] | [Min:Max] |  |
| PRS | -0,214±0,049 | 0,216±0,051 | <0.001 |
|  | [-0,289: -0,073] | [0,121:0,364] |  |
| SVM | -0,073±0,046 | 0,077±0,043 | <0.001 |
|  | [-0,101:0,026] | [-0,020:0,103] |  |
| RF | 0,481±0,154 | 0,495±0,137 | 0.290 |
|  | [0,005:0,550] | [0,005:0,553] |  |
| DL | 0,178±004 | 0,178±0,004 | 0.989 |
|  | [0,162:0,189] | [0,168:0,191] |  |

## DISCUSSION

In general, since the SVM method has lower standard errors and narrower confidence intervals, it can be used as the powerful alternative to the PRS method to calculate PRS only in the control or only the patient group. In previous similar studies, it was seen that only the risk score could be determined for the control group [23]. However, it should not be ignored that other methods can be used if it is desired to determine the PRS of only the control or only the case group by considering them as two different populations. The mean results obtained may have a normal distribution curve for control and case populations to be independent and different from each other. The main reason for this situation should be considered that it is not possible to explain all genetic variation with SNPs, and clinical findings as well as environment-gene interaction should not be ignored. At the highest number of SNPs and highest sample size in the simulation scenario, the results of the classical method and DL methods exceeded 1.00 for both cases and controls, while the SVM and RF methods yielded results in the range of [0.00:0,20]. In the literature, for diseases such as type-II diabetes; there are studies concluding that clinical risk score and PRS are used together, thus increasing the power of case-control separation in the population [24, 25]. In a PRS calculation study using ML in the field of neurology, it was concluded that it is suggested to use SNPs by weighting with ML methods [26]. It has been observed that PRS calculated by MR methods can achieve more successful results in terms of case-control distinction in terms of means compared to the classical method, with different sample sizes and different SNP numbers. In the low number of SNPs, using DL may give better results, and it can be observed that it loses its power as the number of SNPs increases, and it has been seen in previous studies that DL is not strong enough for classification ability is not strong enough for binary GWAS data [27]. Considered to all the PRS calculations methods are based on the weights*SNPs, the part of major alleles which is *SNPs part is all the same. Therefore, source of the difference case and controls is coming from the calculated weights. The weights by calculating DL are having more homogeneity which is not able to separate case and controls can be said. For the weight vectors obtained by the SVM method, the difference between patient and control is significantly high. Because the weight vectors are calculated in such a way that they are farthest from each other. There is a different situation in the RF method. By evaluating the contribution of SNPs to the model, weight vectors were obtained. Considering the normal distribution curve for PRS, it would be a correct approach to compare the methods among themselves with different sample sizes and different SNP numbers, and to compare them with each other. The classical method average can be positive in some cases and negative in some cases. The logarithm of the odds ratio is taken as the weighting method. In this case, too many individuals with a low odds ratio will have negative results. This may be because, in general, the weighted alleles in the controls have lower odds ratios than the weighted alleles in the patients. Although DL’s inability to separation patient-control seems to be a disadvantage at first, if it has a value as a prediction of the score to be calculated or if there is a known risk prediction from previous studies, it can be said that DS method is still a useful method. Because cases and controls groups can take into consideration as a different population group. In similar studies carried out recently, it has been seen that DL weights are used and as a result, results with a wider range of variation are obtained compared to the classical method [28]. To calculate PRS for prostate cancer, 1725 cases 541 controls from Johns Hopkins University (JHU) hospital, 448 cases-1229 controls from Ambry Genetics (AG) company, and 384 controls from NorthShore University (NU) genomics department were selected and 72 with the classical method (PRS). In a study calculated using SNP, in JHU data it was 1.05±0.73 for controls (mean ± std. deviation), 1.41±1.04 for cases, and 1.02±0 for controls in AG firm data. 80 was calculated as 1.46 ± 1.37 for cases, while it was calculated as 0.96 ± 0.63 for controls in the (NU) data. The simulations closest to these results are for the JHU data, it was thought that the results obtained with DL could be 1000 patients-1000 controls 5000 SNPs. In the DL method, it is within the range of [−0.043:1,457] for patients (Table 1). There is a similar situation for the controls, which are located within the range of [0.022:1,432]. Considering the simulation results of 1000 cases and 1000 controls, it can be said that the results obtained with the DL method are the closest to the AR firm data. Considering only the control group for NU data, it was observed that for 500 controls and 5000 SNPs, a result falling between the classical method calculation[-0.645:0.916] and DL [-0.098:1,247] value ranges was obtained. This means that the DL method proof that may be suitable for use in low SNP counts. In another study, emphasizing the multi-ethnic similarity of the polygenic risk score, the PRS was calculated for the breast cancer risk of European and Asian women using 289 SNPs [29]. In this study, 5129 patients 5285 controls were used, and the polygenic risk was calculated as 0.44±0.65 in the case group and 0.12±0.59 in the control group. The closest results to the results of this similar study are the results obtained with 2000 cases 2000 controls 5000 TNP and SVM method; in the case group [-0.152:0.773], in the control group [-0.384:0.757]. If the DL method is to be examined, the results obtained; [0.126:2.155] in the case group and [-0.076:2,150] in the control group (Table 3). In this case, 2000 cases 2000 controls, N=4000 sample size, SVM method can be capable of effectively separating between case and control, with a low number of SNPs like the DL method. PRS should not be confused with diagnostic tests. With PRS, qualitative information about whether individuals have or will have the disease cannot be obtained. However, with PRS it is possible to measure the risk of a progressive disease in individuals [30]. The average risk score to be obtained with the method to be used and the standard deviation values of the scores to be obtained accordingly may be advantageous for the use of PRS, as the range of variation is small. It can be said that ML methods give better results than DL and traditional methods. When calculating PRS with the classical method calculation, it is frequently used to use two different data sets, training, and test sets. Although this is not always easy, it brings a disadvantage for the classical method [31]. By using a single data set, ML and DL methods both avoid the problem of overfitting and can be used as a good alternative method, despite the disadvantage of using two different data sets of the classical method. Previous studies of similar nature have also shown that using GWAS summary statistics in continuous data, ML methods yield better results than the classical polygenic risk score calculation method [32, 33]. In a similar study examining gene-gene and gene-environment interactions related to type-II diabetes, it was observed that the clinical risk score was highly similar to PRS. In another study conducted with similar logic, PRS was used with variables that pose clinical risk [34].

